# Eluding anemone nematocysts: are clownfish deprived of N-acetylated sugars in their skin mucus?

**DOI:** 10.1101/2024.05.11.591997

**Authors:** Sara Heim, Tony Teav, Fabio Cortesi, Hector Gallart-Ayala, Julijana Ivanisevic, Nicolas Salamin

**Affiliations:** Department of Computational Biology, Faculty of Biology and Medicine, University of Lausanne, Lausanne, Switzerland; Metabolomics and Lipidomics Unit, Faculty of Biology and Medicine, University of Lausanne, Lausanne, Switzerland; School of the Environment and Queensland Brain Institute, University of Queensland, Brisbane, Queensland, Australia

**Keywords:** clownfish, damselfish, mucus, N-acetylated-sugars, mutualism, metabolomics

## Abstract

The clownfish - sea anemone system is a great example of symbiotic mutualism where host « toxicity » does not impact its symbiont partner, although the underlying protection mechanism remains unclear. The regulation of nematocyst discharge in cnidarians involves N-acetylated sugars like sialic acid, that bind chemoreceptors on the tentacles of sea anemones, leading to the release of stings. It has been suggested that clownfish could be deprived of sialic acid on their skin surface, sparing them from being stung and facilitating mutualism with sea anemones. In this study, we sampled the skin mucus of two anemone symbionts, the clownfish *Amphiprion akindynos* and the juvenile damselfish *Dascyllus trimaculatus*, as well as two non-symbiotic adult damselfish *Pomacentrus moluccensis* and *P. pavo.* The free and total sialic acid content, including its conjugated form, and three other intermediates of this pathway were quantified using a stable isotope dilution mass spectrometry approach. We found significantly higher levels of sialic acid and its precursor in the non-symbiotic damselfishes. Concentrations of total sialic acid in anemone symbionts ranged between 13 µM and 16 µM, whereas the non-symbiotic damselfishes ranged between 21 µM and 30 µM. The presence of this metabolite and its precursors, as triggers of nematocyst discharge, in anemone symbionts, suggests that this is not the direct mechanism of protection or that the trigger is concentration dependent. This experiment demonstrates that anemone symbionts are not spared by nematocysts because of a lack of N-acetylated sugars, as previously thought.

## Introduction

Mutualistic interactions, implicating species benefiting from close associations with one another, play a fundamental role in shaping biological communities and the evolution of the species involved. In particular, mutualisms can foster species diversification through various mechanisms, one of which involves key innovations that expand the availability of new ecological niches^1,2^. Several examples of species diversification associated with mutualistic interactions have been described^3–5^, but few involve marine species. A notable exception is the mutualistic association of clownfish (Pomacentridae, Amphiprioninae) with giant sea anemones (Actiniaria, Anothozoa), which have become an iconic example in the marine world^6^. This system has been studied in detail to understand the evolutionary and ecological processes that allowed the clownfish clade to diversify in 28 different species through adaptive radiation^7,8^ and has become an attractive model system for diverse biological studies ^9^. An unanswered question regarding the clownfish-sea anemone interaction is how a venomous host can associate so tightly with a potential ‘fish prey’ as a life-long partnership, rendering the system even more intriguing. It is likely that chemical cues are exchanged between these two organisms, providing clownfish with the unique ability to be recognized and subsequently inhabit sea anemones without being harmed, unlike all other susceptible fish species^10,11^. However, the precise molecular pathways involved remain to be elucidated.

Most marine organisms are covered by a mucus layer that possesses important biological and ecological functions and is also the main ‘platform of exchanges’ with the surrounding external environment^12^. Skin surface mucus is likely the primary vector conveying chemical information between clownfish individuals and sea anemones^9^, leading to two main hypotheses underlying this mutualism. The first is chemical camouflage, whereby clownfish can uptake/acquire certain elements from the anemone mucus ^13^. For instance, an exchange in skin surface microbial communities was demonstrated between the two symbiotic partners^14^. The second hypothesis suggests that clownfish skin mucus either contains specific small molecule metabolites or macromolecules that may prevent the discharge of the anemone’s tentacles, or it is deprived of molecular cues present in other fishes, that trigger nematocysts located in the tentacles^15–17^. Recently, the metabolite and lipid profiles of the skin mucus of six clownfish species and two closely related damselfish (Pomacentridae) species have been described^18^, revealing that clownfish mucus is much richer in the sphingolipid class of ceramides compared to damselfish mucus. Nevertheless, it remains unclear whether these sphingolipids play a role in this symbiotic mutualism, potentially conferring clownfish a unique protection from their venomous host.

A good starting point to unravel the mechanisms that allow clownfishes to live within an otherwise toxic environment is to look at the molecular mechanisms that trigger the firing of sea anemone nematocysts^19,20^. Sea anemones are venomous marine invertebrates located across all oceans. Some anemone species can host different species of fish and crustaceans that benefit from this noxious environment as a shelter from predators^21,22^. Cnidarian toxicity is stored in powerful subcellular weapons known as cnidocytes (or cnidae), which are membrane enclosed organelles^23^. The discharge of the nematocysts, located inside the cnidocytes, is fine-tuned by two classes of receptors: mechanoreceptors (hair bundles) and chemoreceptors^24^. The latter binds N-acetylated sugars and the former binds amino acids such as proline^19,20^. N-acetylated sugars are present in mucins, glycoproteins and glycolipids on the skin surface of other organisms (potential preys) and induce structural modifications that lead to the firing of the nematocyst^25^, penetrating the skin of the prey and releasing toxic compounds such as neurotoxins, actinoporins, cytotoxins or other peptides^26^. Numerous studies have shown that N-acetylated sugars, particularly N-acetyl-neuraminic acid (sialic acid), trigger the molecular cascade of nematocyst discharge in sea anemone tentacles^27–30^. The presence of glycosylating agents and nucleotide sugars (UDP-glucose, UDP-N-acetylglucosamine/galactosamine, UDP-D-galacturonic) on surface membranes also plays an important role in establishing symbiotic interactions, such as the recognition and acceptance of dinoflagellate symbionts in sea anemones^31^. In clownfishes, two genes have been identified for their role in the adaptation to host anemones: the Versican core protein and the protein O-GlcNAse^32^, the latter being involved in the cleavage of N-acetylated sugars from macromolecules^33^.

Based on these findings and the hypotheses behind the clownfish-anemone mutualism, we tested for the presence and abundance in clownfish and damselfish mucus of four molecules of the sialic acid pathway that are directly and indirectly involved in protein glycosylation^34,35^: N-acetylneuraminic acid (NeuNAc), N-acetyl-D-mannosamine/hexosamines (ManNAc/HexNAc), Uridine diphosphate N-acetylglucosamine (UDP-GlcNAc) and Cytidine monophosphate N-acetylneuraminic acid (CMP-NeuNAc). We focused on investigating potential differences in symbiotic and non-symbiotic partners of host sea anemones. Glycosylation and glycosylating agents have been studied in *Danio rerio* before^36^, however this field remains understudied in non-model organisms such as Pomacentridae and could be one pilar supporting the hypothesis that clownfishes either have or lack specific compounds enabling them to establish the mutualism with toxic sea anemones^37^. We measured both free and total fractions, including the conjugated form, of NeuNAc, HexNAc, UDP-GlcNAc and CMP-NeuNAc extracted from the skin mucus of the clownfish generalist *Amphiprion akindynos*, the juvenile damsel *Dascyllus trimaculatus* and the adult damselfishes *Pomacentrus moluccensis* and *P. pavo*. These metabolites were measured using a targeted liquid chromatography - tandem mass spectrometry (LC-MS/MS) approach in a dynamic multiple reaction monitoring (MRM) mode. Contrasting both a clownfish to three damselfish as well as anemone symbionts to non-symbiotic damsels allows us to test if the pathway is specific to clownfish and potentially linked to their radiation or rather to the host association itself.

## Results

We quantified the concentration of N-acetylated sugars present in Pomacentridae skin mucus. These molecules are major components of glycoproteins present on the skin surface of many organisms and have been shown to be signaling molecules involved in triggering of nematocyst discharge of sea anemones.

### NeuNAc and its precursors are present in clownfish and damselfish mucus

The metabolites N-acetyl-neuraminic acid (NeuNAc), N-acetyl-hexosamine (HexNAc), UDP-N-acetyl-glucosamine (UDP-GlcNac) and CMP-N-acetyl-neuraminic acid (CMP-NeuNAc) extracted from the skin mucus were detected in all the individuals of *A. akindynos*, *D. trimaculatus, P. moluccensis* and *P. pavo*. Figure 1a and Figure 1b show the extracted ion chromatograms (EICs) of the multiple precursor – product ion (m/z ratio) pairs, specific of NeuNAc and HexNAc (and UDP- and CMP-carrier units) fragmentation patterns. These specific signals measured with state-of-art liquid chromatography-tandem mass spectrometry (LC-MS/MS) confirm that these glycosylating agents are present in both anemone symbionts (*A. akindynos* and juvenile *D. trimaculatus*) as well as the non-symbiotic damselfishes (*P. moluccensis* and *P. pavo)*. The signal specificity demonstrates the presence of NeuNAc and HexNAc.

**Figure 1.**
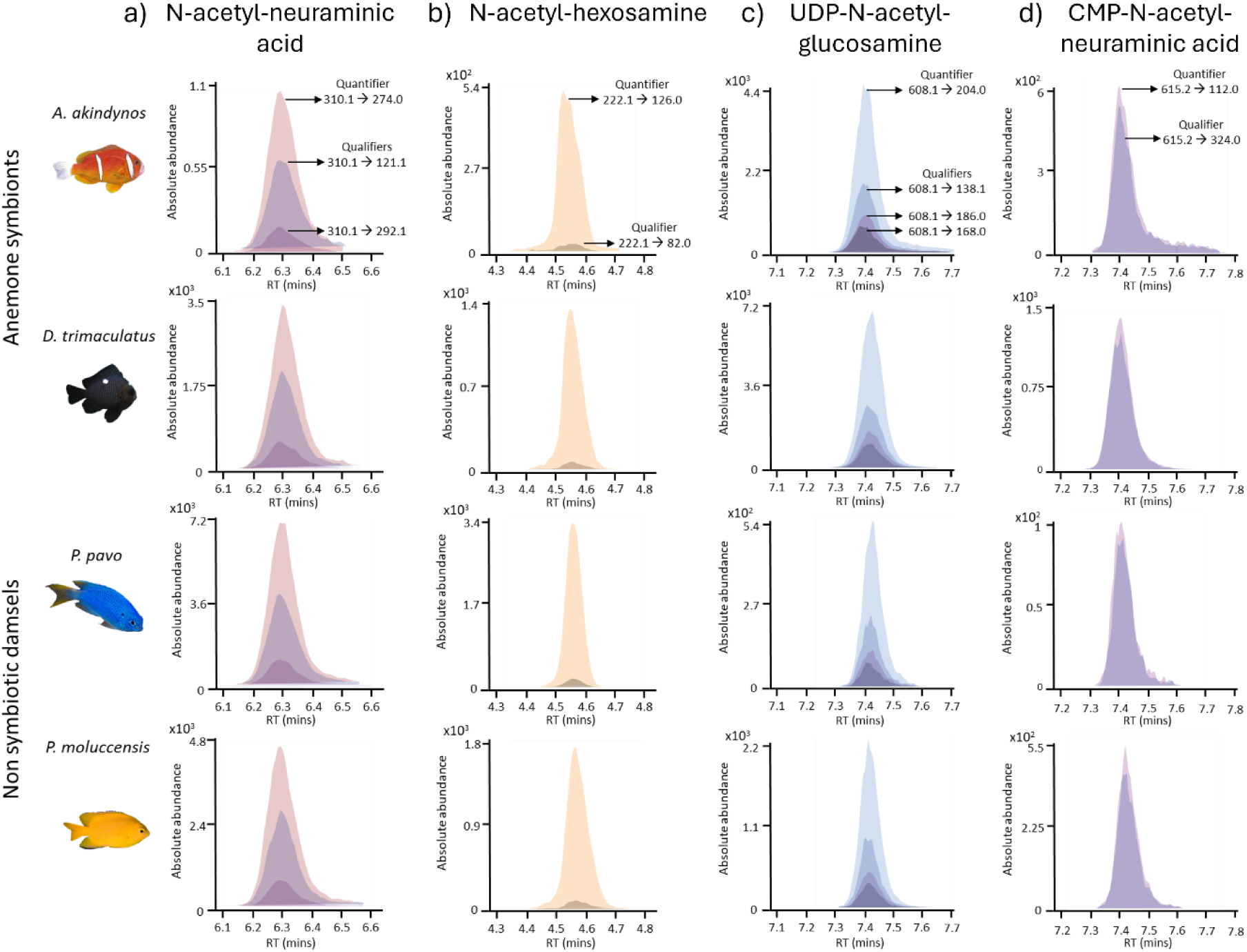
Representative extracted ion chromatograms of the transitions measured in dynamic MRM for (a) N-acetyl-neuraminic acid, (b) N-acetyl-hexosamine, (c) UDP-N-acetyl-glucosamine and (d) CMP-N-acetyl-neuraminic acid in four different species of coral reef fish. The chromatogram was extracted from one sample of each specimen. The retention time (minutes) of the compounds is represented on the abscissa and the abundance of the peaks (total ion counts) is represented on the ordinate. The quantifiers and corresponding qualifiers are indicated on the first chromatograms and applied to all the others.

UDP-GlcNAc and CMP-NeuNAc could not be detected in the total fraction of the mucus extract because they were most likely chemically degraded by the presence of sulfuric acid used for the extraction of total glycosylating agents^39^. Nevertheless, these two metabolites were measured in the free fraction of the polar extract, and they were also present in all four fish species (Figures 1c and 1d). UDP-GlcNAc and CMP-NeuNAc eluted very closely to each other around 7.3 - 7.4 min but could be discriminated by their specific *m/z* of the quantifier and qualifier ion pairs. The ratios of the quantifiers/qualifiers were consistent across both the standards and the endogenous metabolites measured in the samples (See Supplementary table 2 and 3), further confirming the presence of N-acetylated sugars.

### Varying levels of sialic acid and its precursors between clown- and damsel-fishes

We tested if there was a difference in the amount of NeuNAc and ManNAc/HexNAc between the fish species that either interact with sea anemones (‘anemone symbionts’, *A. akindynos* and *D. trimaculatus*) or are potential prey (‘non-symbiotic damsels’, *P. moluccensis* and *P. pavo*). For both total and conjugated amounts, the non-symbiotic damsels had significantly more NeuNAc than anemone symbionts (Figures 2a and 2c, total NeuNAc SE = 4.750, t(16) = −2.185, p-value = 0.044 and conjugated NeuNAc SE = 4.740, t(16) = −2.253, p-value = 0.039). The mean values of *P. moluccensis* and *P. pavo,* which do not interact with sea anemones, are nearly two times larger for total NeuNAc than the other fish species (∼26 µM versus 14.5 µM). When comparing the concentration of NeuNAc between the clownfish *A. akindynos* and the other 3 damselfish species, there was no significant difference in the total nor in the conjugated amounts (Figure 2b and 2d).

**Figure 2.**
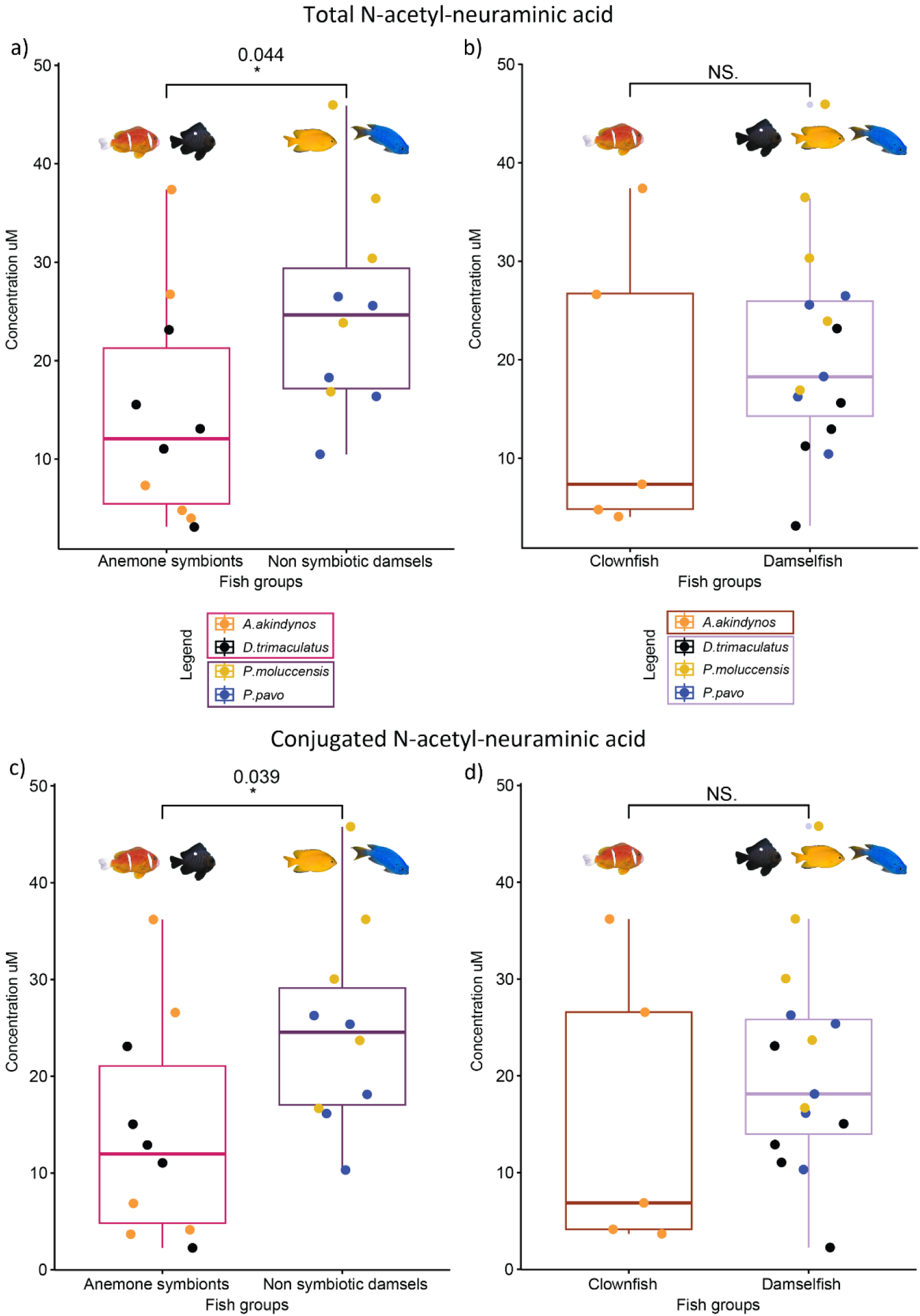
Boxplots of total (a) and (b) and conjugated (c) and (d) amounts of N-acetyl-neuraminic acid measured in the fish species of this study (n = 5 individuals per species). The p-values above the boxplots were calculated using the anova test and contrasting anemone symbiotic fishes and non-symbiotic damselfishes, as well as the clownfish, A. akindynos, to the three other damselfish species. A. akindynos and D. trimaculatus are grouped as anemone symbionts whereas P. moluccensis and P. pavo as non-symbiotic damsels. The horizontal line in the boxplots represents the median concentration of NeuNAc and the whiskers delimit the upper and lower quartiles. The grey dots neighboring a colored dot beyond the whiskers represent the outliers.

The total and conjugated amounts of N-acetyl-hexosamines were also significantly lower in anemone symbionts compared to the non-symbiotic *P. moluccensis* and *P.pavo* (Figures 3a and 3c, total HexNAc SE = 7.120, t(16) = −2.399, p-value = 0.029 and conjugated HexNAc SE = 6.940, t(16) = −2.407, p-value = 0.028). The clownfish also had less total and conjugated HexNAc when comparing these concentrations to the ones of the other 3 damselfish species (Figures 3b and 3d, total HexNAc SE = 8.220, t(16) = −2.1125, p-value = 0.049 and conjugated HexNAc SE = 8.010, t(16) = −2.140, p-value = 0.048). We also measured the difference of total, free and conjugated NeuNAc and HexNAc between the 4 species (Supplemenary Figures 1 and 2). Total and conjugated amounts of NeuNAc were found to be the most abundant in the damselfish *P. moluccensis* (Supplementary Figures 1a and 1c). Free NeuNAc content was significantly higher in the anemone symbionts compared to the non-symbiotic damsels (Supplementary Figure 1b, SE = 0.123, t(16) = 2.480, p-value = 0.025), although intraspecies variability was high in anemone symbionts. On the other hand, total and conjugated HexNAc were most abundant in the blue damselfish *P. pavo* (Supplementary Figures 2a and 2c), with concentrations ∼ 53 µM whereas the clownfish *A. akindynos* had the lowest amounts (∼18 µM). There was no significant difference in the measurements of free HexNAc among the fish species (Supplementary Figure 2b)

**Figure 3.**
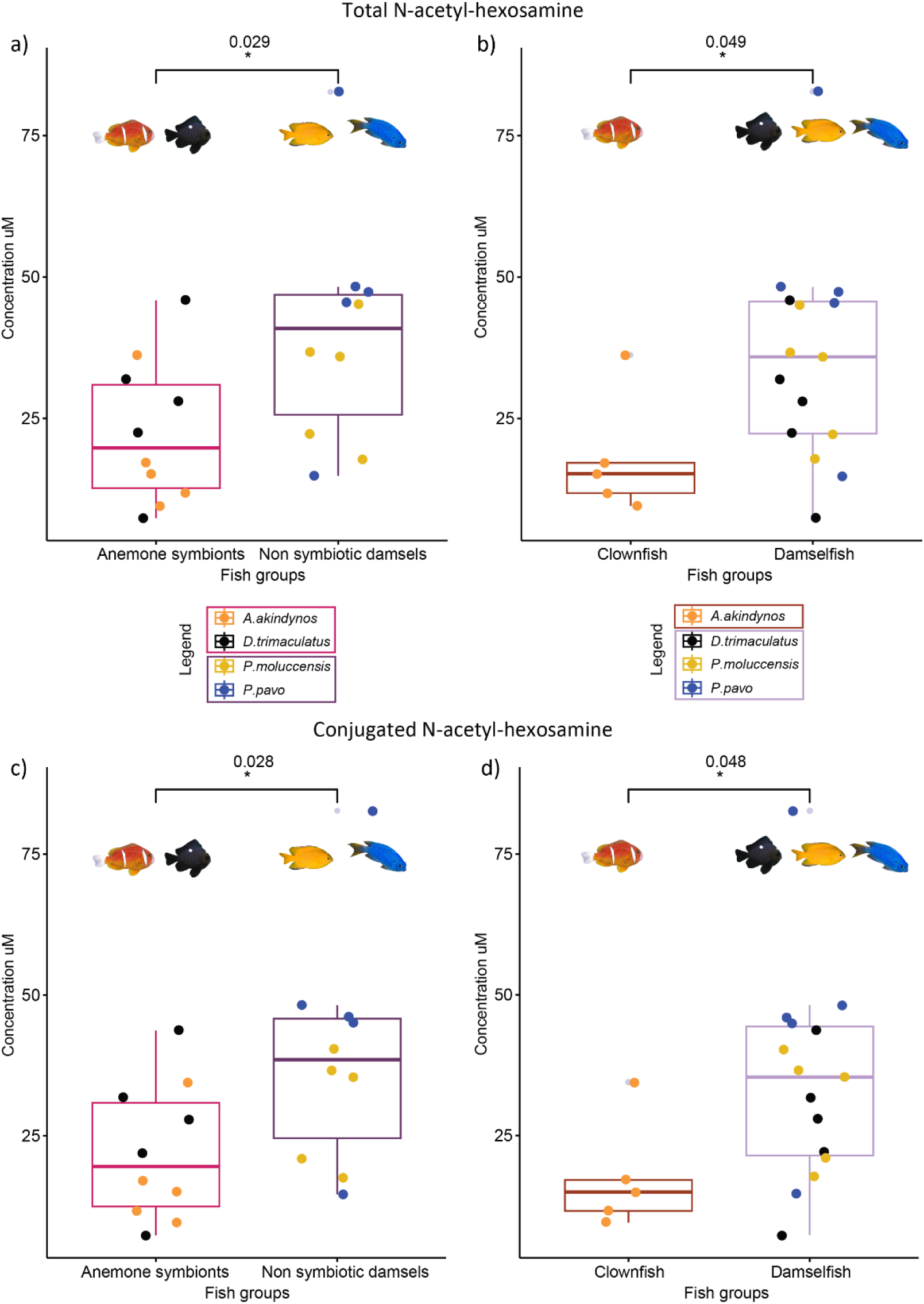
Boxplots of total (a) and (b) and conjugated (c) and (d) amounts of N-acetyl-hexosamine measured in the fish species of this study (n = 5 individuals per species). The p-values above the boxplots were calculated using the anova test and contrasting anemone symbiotic fishes and non-symbiotic damselfishes, as well as the clownfish, A. akindynos, to the three other damselfish species. A. akindynos and D. trimaculatus are grouped as anemone symbionts whereas P. moluccensis and P. pavo as non-symbiotic damsels. The horizontal line in the boxplots represents the median concentration of HexNAc and the whiskers delimit the upper and lower quartiles. The grey dots neighboring a colored dot beyond the whiskers represent the outliers.

We also tested potential differences in the carrier units of the sialic acid pathway. UDP-GlcNAc and CMP-NeuNAc were present at very low concentrations (< 2uM) in fish mucus extracts: between 7×10^-6^ and 1.6 µM for UDP-GlcNAc and between 3×10^-4^ and 0.2 µM for CMP-NeuNAc (Supplementary figure 3). When comparing these two metabolites between anemone symbionts and coral associated fish and between clownfish and damselfish species, we found no significant differences (Supplementary Figure 3).

## Discussion

In this study, we report the concentrations of four metabolites of the N-acetyl-neuraminic acid metabolic pathway (NeuNAc/sialid acid)^40^ in the mucus of four damselfish species; two anemone-associated species, the obligate symbiont clownfish, *A. akindynos* and the partial symbiont, *D. trimaculatus*, which only interacts during the juvenile stage, and two non-symbiotic species, *P. moluccensis* and *P. pavo*. We assumed that the pathway is important in the context of host-anemone mutualism because it is involved in protein and lipid glycosylation in fish skin mucus^36^. Based on extensive work which showed that NeuNAc is a trigger of cnidarian nematocysts^20,41–43^, we hypothesize that this small molecule metabolite may be absent in anemone symbiotic partners such as clownfishes or the ‘partially symbiotic’ juvenile *D. trimaculatus*. To the best of our knowledge, only one study that had previously measured N-acetylneuraminic acid in the skin mucus of *A. ocellaris* and two other fish that do not interact with sea anemones (the moonwrasse and the scissortail sergeant) using HPLC-FLD^44^, showing that the clownfish had 40 times less NeuNAc compared to the other two fish species tested. However, the presence and concentration of HexNAc and the carriers UDP-GlcNAc and CMP-NeuNAc were unknown. Further, it is important to compare closely related species showing full, partial and no interaction with sea anemone to better assess whether clownfish ability to live within sea anemone is due to the presence or absence of these metabolites.

Using a targeted liquid chromatography - tandem mass spectrometry (LC-MS/MS) approach, we show that anemone symbionts have less NeuNAc and HexNAc compared to other damselfishes but the difference is less striking than previously shown. The discrepancy might be due to two main factors: first, our study used a stable isotope dilution approach for the internal calibration of NeuNAc and HexNAc; second an external calibration was applied for UDP-GlcNAc and CMP-NeuNAc. Using the internal standard response, we are able to translate the peak areas to concentrations of the targeted metabolites and correct for potential differences in matrix effects in the mucus collected from different species^45^. The metabolites are separated on a hydrophilic liquid chromatographic column and measured by multiple reaction monitoring mass spectrometry on a triple quadrupole, allowing for the higher selectivity and higher specificity for the molecules of interest compared to an HPLC-FLD system^46^. This latter technique is less specific, and the chromatograms obtained in previous studies from the clownfish samples show an important matrix effect, making the peak integration tedious. From our data, we observe that the skin mucus of the clownfish *A. akindynos* and the juvenile *D. trimaculatus,* which live among host anemones without being stung, also contains NeuNAc (half the amount measured in the non-symbiotic damselfishes) and its precursors. Both species had similar amounts of NeuNAc, although the clownfish showed a high variability among individuals (Supplementary figure 2). On the other hand, the concentration of total and conjugated HexNAcs was at its lowest in the clownfish *A. akindynos* (∼ 17 µM) with much less variability. The ‘facultative symbiotic’ juvenile *D. trimaculatus* presents total and conjugated HexNAcs amounts similar to those of the non-symbiotic *P. moluccensis* (∼ 27 µM and 31 µM respectively), suggesting that the sugar class in the skin mucus varies across damselfish species. A comparison of more clownfishes and other damselfish species is needed to test this hypothesis conclusively.

Although there is evidence that these metabolites can bind to chemoreceptors on sea anemone tentacles and trigger the release of the nematocyst^25^, their presence in the mucus of anemone symbionts shown in our study together with previous genomic evidence of genes coding for O-GlcNAcase and Versican core protein under positive selection in clownfishes^47^, indicate that the molecular pathway behind this mutualism is more complex. These metabolites are often found on the terminal end of glycans that could play an important role in recognizing mutualistic partners^48,49^.

N-acetylated sugars and other sugar molecules (monosaccharides and glycans) commonly serve as glycosylating agents to form glycoproteins and glycolipids^50^. These macromolecules are often located on the surface of cell membranes and are involved in different biological processes, such as cell-cell communication or cell-microbe interactions ^51^. Glycans bind on specific atoms of certain amino acids giving rise to N-linked glycans when they attach to the nitrogen atom of Asparagine and O-linked glycans when they attach to the oxygen atom of Serine or Threonine residues^52^. These long chains of sugars can be analyzed by MS and MS^n^ to obtain fragment ions and deduce the sugar composition of the branch. However, these technologies cannot discriminate between different isoforms with the same mass values ^53^. It is also challenging to separate the isoforms using a chromatographic system as they have extremely similar chemical properties and often co-elute. For this reason, in our study, we refer to N-acetyl-mannosamine as N-acetyl-hexosylamine. Even when using the internal standard of N-acetyl-mannosamine, the number of ions measured with the same m/z could be rising from other isomeric sugars (N-acetyl-galactosamine or N-acetyl-glucosamine). Recent evidence supporting the hypothesis that larger sugar chains (glycans) play a role in this mutualism was demonstrated in a study monitoring clownfish mucus glycan composition before and after several weeks of established mutualism. Glycan profiles in the clownfish *A. percula* mucus significantly changed after three weeks of contact with the host, but these profiles were lost when the clownfish was removed from the host^54^. Unfortunately, the identification of the glycans was not carried out, therefore it remains unknown which sugar chains could be playing important roles between clownfishes and their hosts. Moreover, the relatively ‘long’ period needed to observe changes in glycan composition and their loss within such a short time could be due to the exchange in microbial communities between the two organisms^53^. Despite all the results obtained by the clownfish scientific community, a big gap remains in understanding the metabolic pathways responsible for this intriguing marine mutualism.

By analyzing small polar metabolites, we were able to detect and quantify the small building blocks of potentially larger molecules such as glycans, glycoproteins or glycolipids present in the mucus of *A. akindynos*, *D. trimaculatus*, *P. pavo* and *P. moluccensis*. We observe a significant difference in the total and conjugated content of NeuNAc and HexNAc among the anemone symbionts and the non-symbiotic damselfishes (Figure 2 and 3). Nevertheless, these differences are less striking than previously reported in the study from 2015 and there was also some variability between the individuals of each species. These results show that both anemone symbionts and non-symbiotic damsels have glycosylating agents on their skin surface mucus that are products of hydrolyzed/deconjugated glycans via the addition of sulfuric acid in our extraction protocol. A recent study also show that glycans in the skin mucus of clownfish undergo fine tuning after several weeks of contact with their host^55^ and this time frame suggests that microbial communities may contribute to this process^54–56^. Many studies have revealed how glycosylation, as a posttranslational modification of proteins, impacts the final 3-D protein folding ^56–58^. In addition, glycans and oligosaccharide chains are flexible in space and can readapt their conformation dynamically ^59,60^. These physical properties lead to the hypothesis that clownfish and other anemone symbionts may possess similar glycoproteins to other non-symbiotic fishes, but the conformation may differ and the glycan building block may be organized differently. For example, damselfishes could have glycoproteins exposing N-acetylated sugars of the oligosaccharide chains that bind to the sea anemone tentacles triggering the nematocyst discharge. On the other hand, clownfish may have glycoproteins that present another 3-D structure whereby the glycosylating agents may be internally folded, therefore not accessible to bind the anemone’s chemoreceptors. This should be further tested using a proteomics approach.

In conclusion, the molecular mechanisms behind this iconic symbiotic mutualism remain unclear and our results lead to several new hypotheses. Throughout the years, this scientific dilemma has been tackled by almost all omics techniques, except by proteomics. Diving into analyzing macromolecules in the damselfish family is the next step to consider for solving the evolutionary and ecological secret behind the clownfish-anemone mutualism. In addition, investigating their conformation in space could also be an exciting approach, although extremely challenging.

## Materials and Methods

### Sampling of fish mucus

Adults from the Barrier Reef anemonefish [*Amphiprion akyndinos* (n=5)], two damselfish species [*Pomacentrus moluccensis* (n=5), *Pomacentrus pavo* (n=5)] and juveniles of the Threespot Dascyllus damselfish [*Dascyllus trimaculatus* (n=5)] were sampled on snorkel from the reefs surrounding Lizard Island (14° 40 S, 145° 27 E), Northern Great Barrier Reef, Australia (Figure 4). All fish were caught on the reef using hand nets and clove oil, brought to the boat to be anesthetized individually in a solution of seawater and Tricaine methanesulfonate (100 mg/l) for several minutes (1–2 min). They were then removed from the solution and placed in a sterilized glass petri dish for mucus collection (3–4 scrapes per side) with the help of a soft cell scraper (SARSTEDT, Nümbrecht Germany) starting from the pectoral fin until the end of the tail, as described in detail in Heim et al. (2023). The mucus was washed off from the cell scraper using 500 μl of UHPLC water, transferred into an Eppendorf tube, kept in ice on the boat and then transferred to liquid nitrogen for storage at the Lizard Island Research station. After manipulation, fish were put into fresh sea water until they awakened and regained mobility, and later returned to the reef/host anemone where they were originally captured. A schematic illustration of the sampling procedure, data acquisition and NeuNAc pathway can be found in Figure 4.

**Figure 4.**
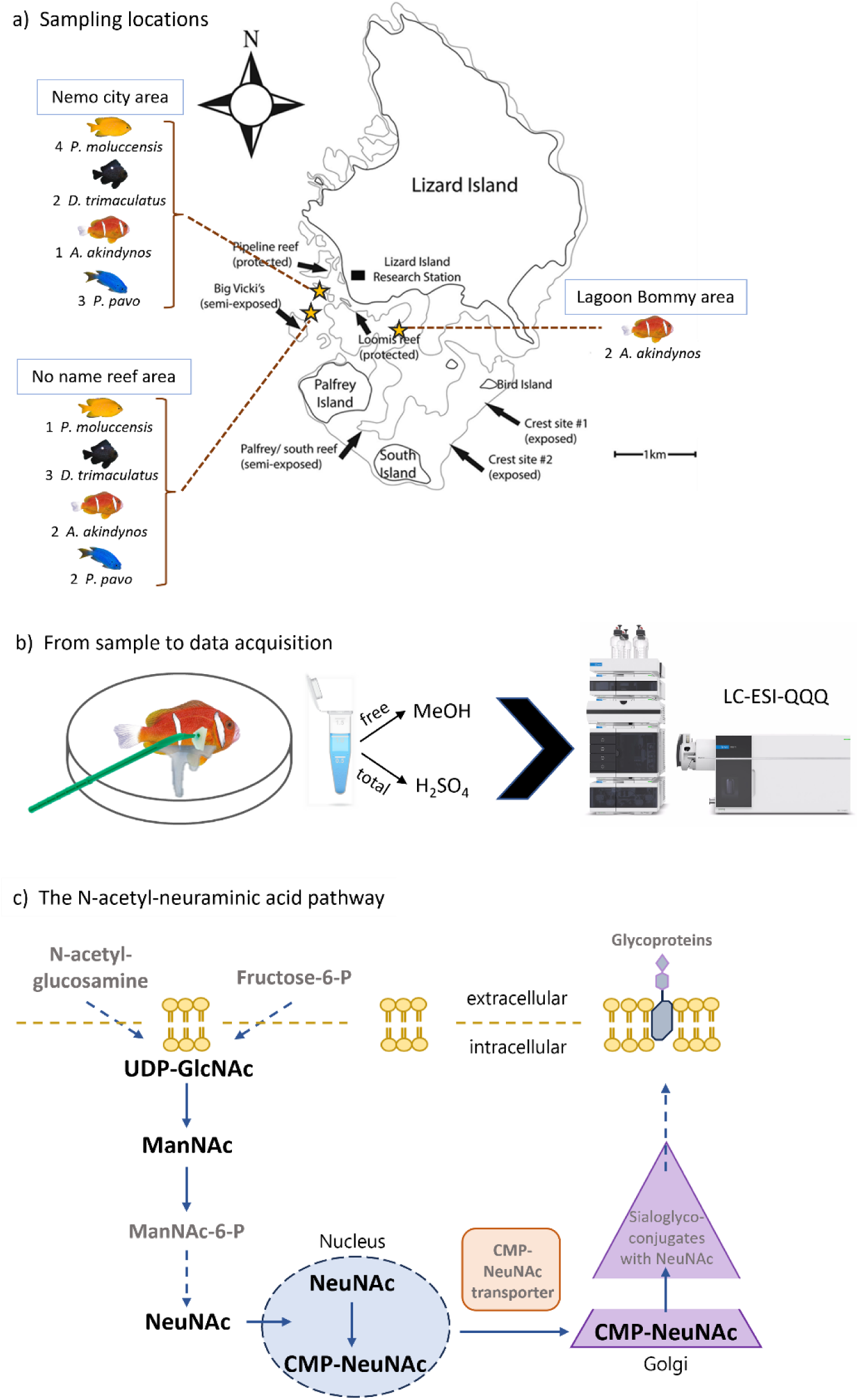
a) Map of Lizard Island in Australia with the main reef areas around the island. The stars indicate the sampling locations of the fish individuals used in this study. The number of individuals per location is indicated around the map. Fish were caught free diving at depths between 1-4 meters. b) Schematic summary of how the mucus was sampled and the solvents used for the extraction of free amounts and total amount of the metabolites targeted and measured using an LC-ESI-QQQ system. c) The N-acetyl-neuraminic acid pathway in vertebrates. In bold are the metabolites analyzed in this experiment and grey are other players of this pathway.

Ethics declaration: All procedures were performed in accordance with the permits from GBRMPA (G17/38160.2) and Animal Ethics (AE000569) from the University of Queensland, Brisbane, Australia, and the Fisheries Act from the Queensland Government (207975). All authors complied to the ARRIVE guidelines. Experimental protocols were reviewed and approved by Native/Exotic Wildlife and Marine Animals AEC (NEWMA) committee of the University of Queensland.

### Reagents and chemicals preparation

For the quantification of the target metabolites, their corresponding chemical standards (STD) and isotopically labelled standards (ISTD) were purchased from Sigma-Aldrich (St. Louis, USA), Toronto Research Chemicals (Toronto, Canada) and Cayman Chemical (Michigan, USA). UHPLC-MS Grade water, acetonitrile, methanol, ammonium formate and formic acid were purchased from Biosolve (Dieuze, France). All standards and internal standards were prepared and diluted in water to obtain a final concentration of 1mM. The stock calibration mixture of standards was initially prepared in water. These stock solutions were then further diluted to match the concentration ranges detected in test samples for the selected metabolites in fish mucus. The curve was designed using the four chemical standards and spinked with the internal standards, covering both a higher range and a lower range of concentrations of the metabolites (test sample concentrations correspond to the mid-range of the curve, between Cal5 and Cal7 points). The concentrations of the ten-point calibration curve can be found in Supplementary Table 1.

Similarly, the stock internal standard (IS) mixture of ^13^C_3_-NeuNAc, ManNAc-d_3_ was prepared in water, with a final concentration of 100 µM.

We reported the concentrations of NeuNAc and ManNAc based on internal calibration (using the spiked ISTD at known concentration) while the concentrations of UDP-GlcNac and CMP-ManNac were calculated based on the external multipoint calibration curve (the corresponding ISTD is not available on the market). The conjugated sugars were calculated based on the total concentration measured minus the concentrations of the free sugars.

### Sample preparation for the extraction of free NeuNAc, ManNAc, UDP-GlcNAc and CMP-NeuNAc

For consistency in the chemical extraction, samples were first transferred into lysis tubes (soft tissue homogenizing CK 14 tubes, Bertin Technologies, Rockville, MD, US), homogenized for 5 seconds with ceramic beads (1.4mm zirconium oxide beads) in a Cryolys Precellys 24 sample homogenizer 5 seconds at 10,000 rpm (Bertin Technologies, Rockville, MD, US) to obtain a homogenous matrix from which metabolites were extracted. The Precellys was air-cooled by the Cryolys at a flow rate set at 110 L/min at 6 bars.

For absolute quantification of NeuNAc, ManNAc, andquantification of UDP-GlcNAc and CMP-NeuNAc (using the external calibration curve), samples were prepared by adding 15 μL of the diluted stock ISTD mixture in methanol to an aliquot of homogenized fish mucus (200 µL). Metabolites were extracted by adding ice-cold MeOH (400 μL). Samples were then vortexed and centrifuged for 15 minutes at 4°C and 21000g and the resulting supernatants were transferred to new Eppendorf tubes and evaporated to dryness with a vacuum concentrator (LabConco, Missouri, US). Dried extracts were re-suspended in HPLC water (100 μL), vortexed, sonicated for 1 min and centrifuged for 15 min at 21000*g* at 4 °C. The resulting supernatants of fish mucus extracts were transferred to LC-MS vials prior to the injection. Ten-point calibration curves were generated following the same procedure as for the samples.

Hazard statements for MeOH: [H225 - H301 + H311 + H331 – H370](https://www.sigmaaldrich.com/IN/en/life-science/safety/hazard-and-precautionary-statements#hazard)

### Extraction of total NeuNAc and ManNAc

An aliquot of homogenized fish mucus samples (25 μL) was placed in LC-MS glass vials, spiked with 15 μL of the diluted stock IS mixture (at 15 μM) and extracted using sulfuric acid (60 μL at 63 mM). Samples were vortexed for 1 minute, incubated at 80 °C for one hour and cooled down at room temperature for 15 min prior to centrifugation for 15 min at 21000*g* at 4 °C. The resulting supernatant was transferred to new LC-MS vials prior to injection.

Hazard statements for sulfuric acid: [H290 – H314](https://www.sigmaaldrich.com/IN/en/life-science/safety/hazard-and-precautionary-statements#hazard)

### Hydrophilic Liquid Chromatography Tandem Mass Spectrometry (HILIC-MS/MS) analysis

Free NeuNAc, ManNAc, UDP-GlcNAc, CMP-NeuNAc and total NeuNAc, ManNAc in fish skin mucus were measured by quantitative stable-isotope dilution assisted assay by hydrophilic interaction liquid chromatography coupled to tandem mass spectrometry (HILIC-MS/MS) in positive ionization mode using a 6495 triple quadrupole system (QqQ) interfaced with 1290 UHPLC system (Agilent Technologies, Santa Clara, US). The chromatographic separation was carried out in a SeQuant^®^ ZIC-pHILIC, 5 μm, 100 mm × 2.1 mm I.D. column (Merck, Darmstadt, Germany). Mobile phases were composed of A = 20mM Ammonium acetate and 20mM Ammonium hydroxide in water and B = acetonitrile. The linear gradient elution from 90% B (0–1.5 min) to 50% B (8–11 min) to 45% B (12– 15 min) was applied. The initial gradient conditions were restored within 1 min and a 9-min post-run equilibration was applied to maintain the system reproducibility. The flow rate was 300 μL min^−1^, column temperature 30°C and the sample injection volume was 2 μl. ESI source conditions were set as follows: dry gas temperature 250 °C, nebulizer 35 psi and flow 15 L/min, sheath gas temperature 400 °C and flow 8 L/min, nozzle voltage 1000 V, and capillary voltage 3000 V. The data were acquired in a dynamic Multiple Reaction Monitoring (dMRM) mode with a total cycle time of 500 ms. Two transitions were used to monitor each compound, and quantifiers *m/z* 310➔274 and *m/z* 222➔126 were used for the quantification of NeuNAc and ManNAc, respectively. The quantifiers for UDP-GlcNAc and CMP-NeuNAc were *m/z* 608.1➔204.0 and *m/z* 615.2➔112.

### Data Processing

Raw LC-MS/MS data were processed using the Agilent Quantitative analysis software (version 10.0 MassHunter, Agilent technologies). For absolute quantification, calibration curves and the stable isotope-labelled internal standards (IS) were used to determine the response factor. Linearity of the calibration curves was evaluated for each metabolite; in addition, peak area integration was manually curated and corrected where necessary. The concentrations of total, free and conjugated (total - free) NeuNAc and ManNAc (N-acetyl-hexosamines) were determined based on the proportion of the corresponding internal standard versus the abundance of endogenous metabolite. The precursor of N-acetyl-neuraminic acid, N-acetyl-mannosamine, is a monosaccharide that can be present in different isomers and various epimers; therefore term ‘N-acetyl-hexosamine (HexNAc)’ is used to relate to monosaccharides co-eluting with N-acetyl-mannosamine such as N-acetyl-glucosamine or N-acetyl-galactosamine.

To evaluate the proportion of each metabolite in the different fish species, we generated a linear model for each metabolite using the lm function in R. We performed a Tukey post-hoc test (TukeyHSD function in R) on the anova for *A. akindynos*, *D. trimaculatus, P. moluccensis* and *P. pavo* to determine if there were statistically significant differences at a p-value of < 0.05, between any pair of species. We also computed the estimated marginal means of the linear model for the contrasts between ‘anemone-symbionts’ (grouping *A. akindynos* and *D. trimaculatus*) and ‘coral-associated’ fishes (grouping *P. moluccensis* and *P. pavo*) as well as between the clownfish and the 3 damselfish of this study, using the emmeans and contrast functions (http://www.rstudio.com/ version 2023.12.0.369). These results were represented in boxplots.

## Acknowledgements

We want to thank Prof. Karen Cheney, Emeritus Prof. Justin Marshall, Abigal Shaughnessy, Jasper Stead and Carl Santiago for helping with the fish catching and sample logistics for this project at the Lizard Island Research Station and the University of Queensland. We would also like to thank the staff at the Lizard Island Research Station for their support during fieldwork and acknowledge the Dingaal, Ngurrumungu and Thanhil peoples as traditional owners of the lands and waters of the Lizard Island region from where the specimens for this study were collected.

## Data availability

Raw data and data tables used for this study are accessible on zenodo via the following link: https://zenodo.org/records/11047437

