## Supplementary Table, Supplementary Figure for "Eluding anemone nematocysts: are clownfish deprived of N-acetylated sugars in their skin mucus?"

**Supplemetary Data**

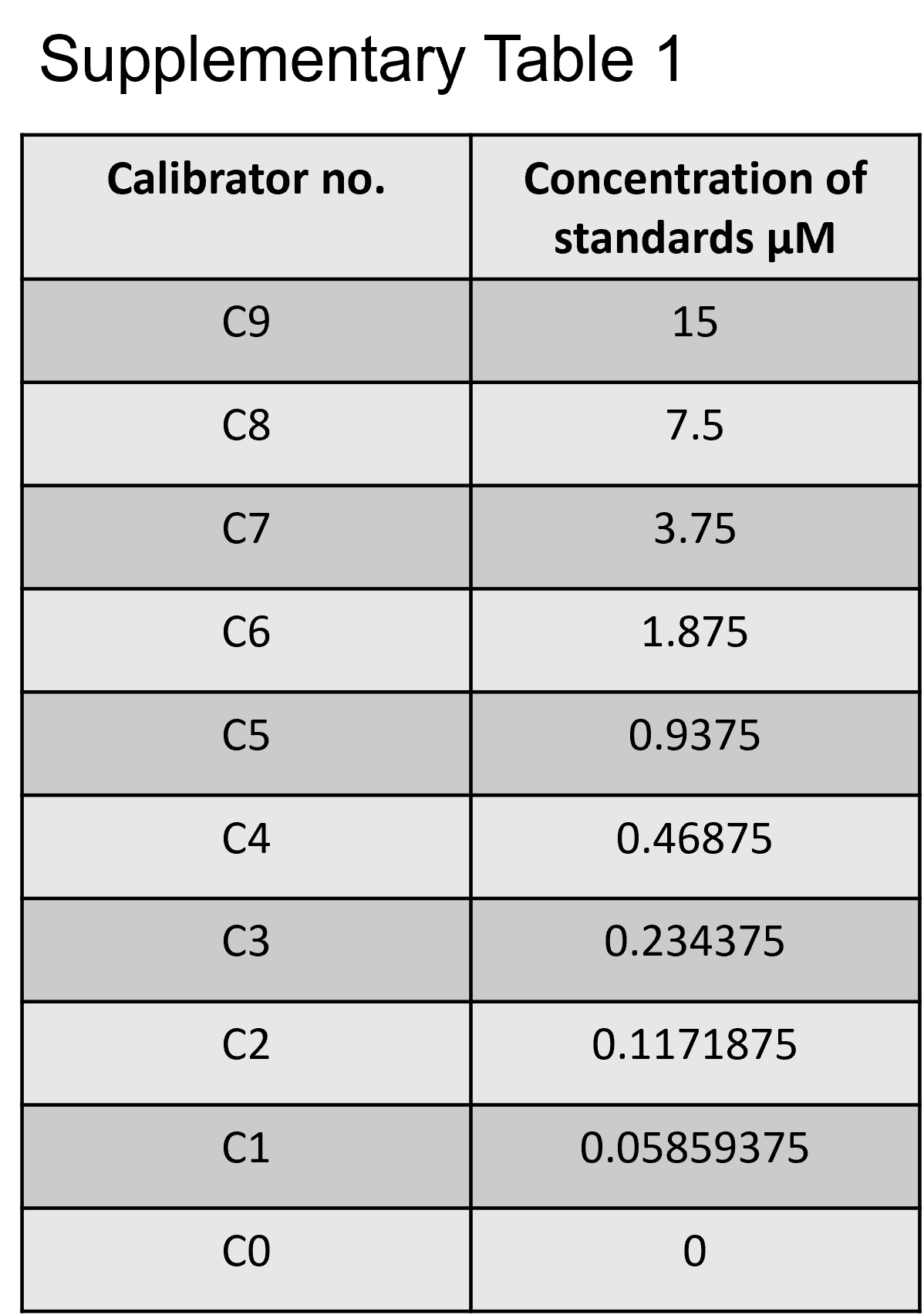
Supplementary table 1 Concentrations in calibrators of NeuNAc, HexNAc, UDP-GlcNAc and CMP-NeuNAc

Supplementary Table 1 The ten-point calibration curve with its corresponding standard mix concentrations. The curve was adapted to the concentrations detected in test samples of clownfish and damselfish mucus. Each C point contains the standard mix of NeuNAc, ManNAc, UDP-GlcNAc and CMP-NeuNAc which was spiked with the internal standard of ^13^C_3_-NeuNAc and ManNAc-d_3_ .

Supplementary table 2 Ratios of quantifiers/qualifiers for NeuNAc and HexNAc

| Calibrator or Species | NeuNAc H+ | | HexNAc H+ | |
| --- | --- | --- | --- | --- |
|  | SD | ratio 274.0/292.1 | SD | ratio 126.0/82.0 |
| Cal6 | 2.28 | 56.6 | 2.48 | 8 |
| Cal7 | 2.28 | 52.9 | 2.48 | 11.5 |
| Cal8 | 2.28 | 53.9 | 2.48 | 6.2 |
| *A. akindynos* | 2.08 | 58.36 | 1.21 | 6.62 |
| *D. trimaculatus* | 2.27 | 54.96 | 2.41 | 6.96 |
| *P. pavo* | 0.64 | 55.9 | 0.78 | 5.96 |
| *P. moluccensis* | 0.93 | 56.02 | 0.74 | 5.64 |

Supplementary Table 2 The ratios of the quantifiers/qualifiers are calculated for all samples per metabolite. Three points of the calibration curve are shown with the mean of the samples per species (n=5). These ratios are consistent throughout the analytical run further confirming the measurements of both NeuNAc and HexNAc.

Supplementary table 3 Ratios of quantifiers/qualifiers for carrier units UDP-GlcNAc and CMP-NeuNAc

| Calibrator or Species | UDP-GlcNAc H+ | | CMP-NeuNAc H+ | |
| --- | --- | --- | --- | --- |
|  | SD | ratio 204.0/186.0 | SD | ratio 324.0/315.4 |
| Cal6 | 0.97 | 18.90 | 2.12 | 87.10 |
| Cal7 | 0.97 | 18.80 | 2.12 | 91.60 |
| Cal8 | 0.97 | 17.90 | 2.12 | 93.70 |
| *A. akindynos* | 2.89 | 19.53 | 7.85 | 86.40 |
| *D. trimaculatus* | 0.90 | 22.44 | 3.63 | 85.74 |
| *P. pavo* | 1.06 | 21.84 | 5.84 | 90.28 |
| *P. moluccensis* | 2.48 | 19.42 | 6.24 | 86.24 |

Supplementary Table 3 The ratios of the quantifiers/qualifiers are calculated for all samples per metabolite. The mid points of the calibration curve are shown with the mean of the samples per species (n=5). These ratios are consistent throughout the analytical run further confirming the measurements of both carrier units UDP-GlcNAc and CMP-NeuNAc.

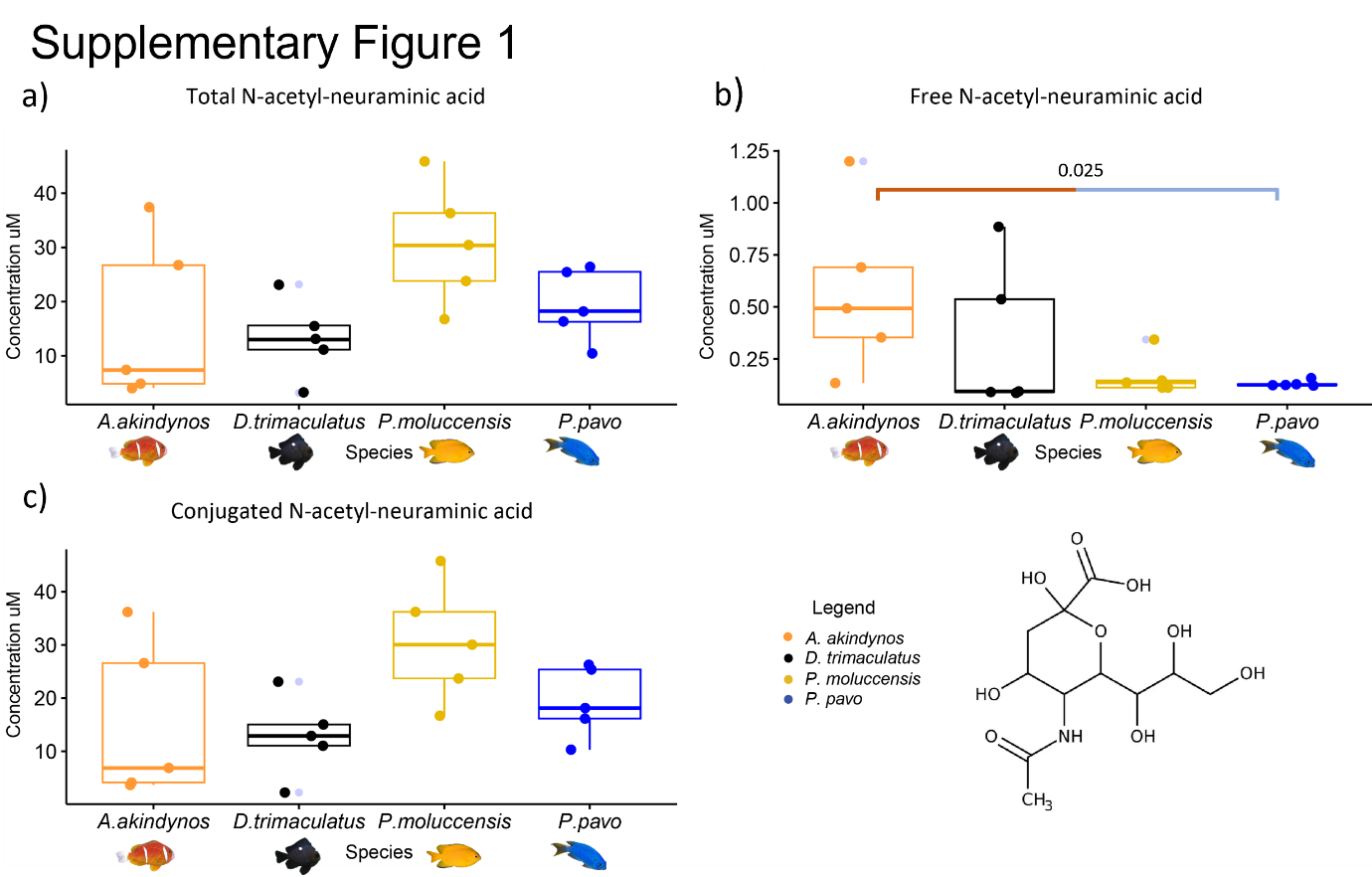

Supplementary figure 1 Levels of N-acetyl-neuraminic acid measured in each species (n=5). In a) are the total amounts of NeuNAc, in b) are the free amounts of NeuNAc and in c) are the conjugated amounts (total-free) represented in boxplots. Free NeuNAc content differences were calculated using the anova test and contrasting anemone symbiotic fishes and non-symbiotic damselfishes. The whiskers represent the upper and lower quartiles whereas the horizontal line is the median concentration within each species. On the right corner is the molecular structure of NeuNAc. The grey dots neighboring a colored dot beyond the whiskers represent the outliers.

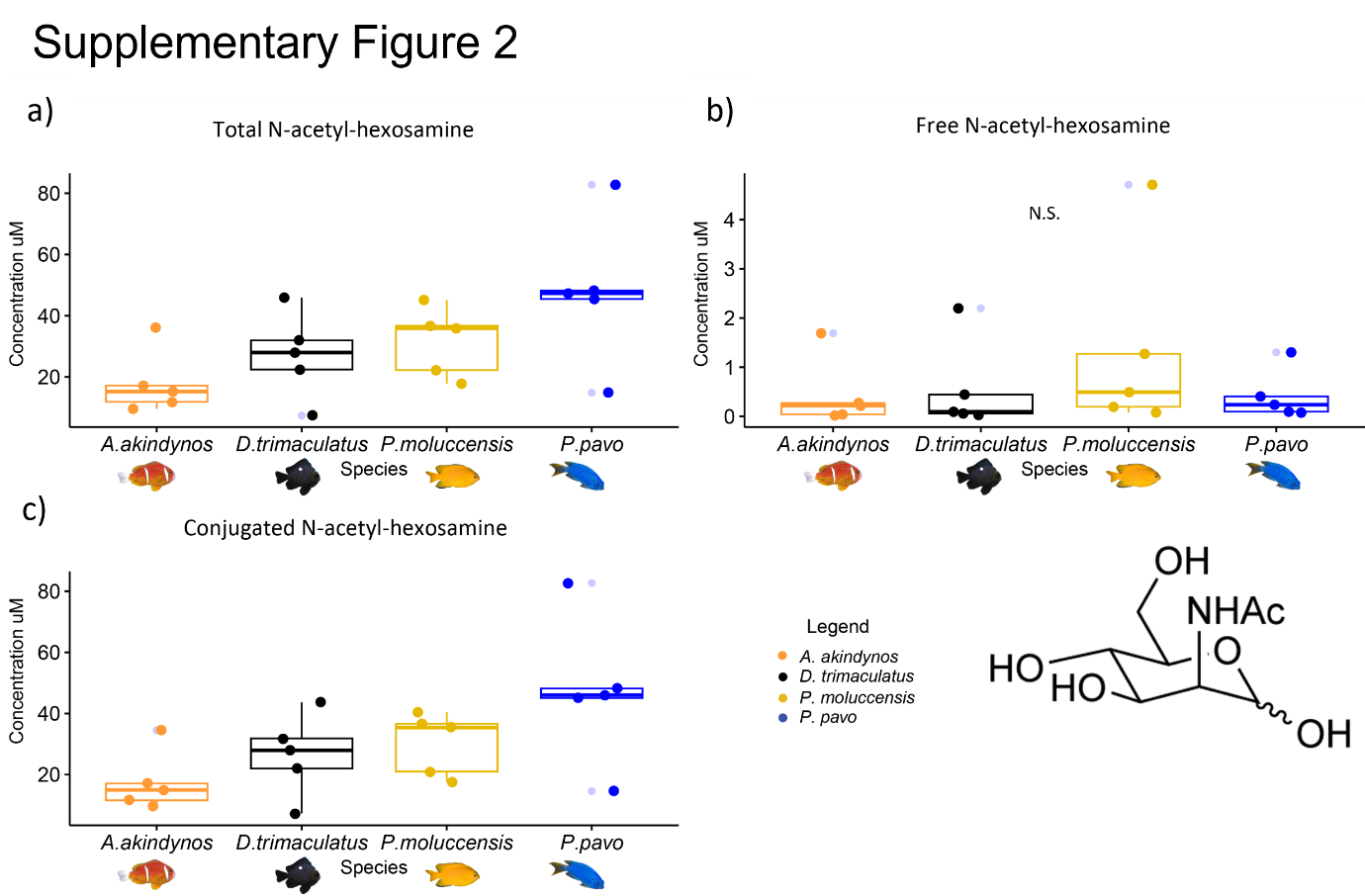

Supplementary figure 2 Levels of N-acetyl-hexosamine measured in each species (n=5). In a) are the total amounts of HexNAc, in b) are the free amounts of HexNAc and in c) are the conjugated amounts (total-free) represented in boxplots. Free HexNAc content differences were calculated using the anova test and contrasting anemone symbiotic fishes and non-symbiotic damselfishes. The whiskers represent the upper and lower quartiles whereas the horizontal line is the median concentration within each species. On the right corner is the molecular structure of HexNAc. The grey dots neighboring a colored dot beyond the whiskers represent the outliers.

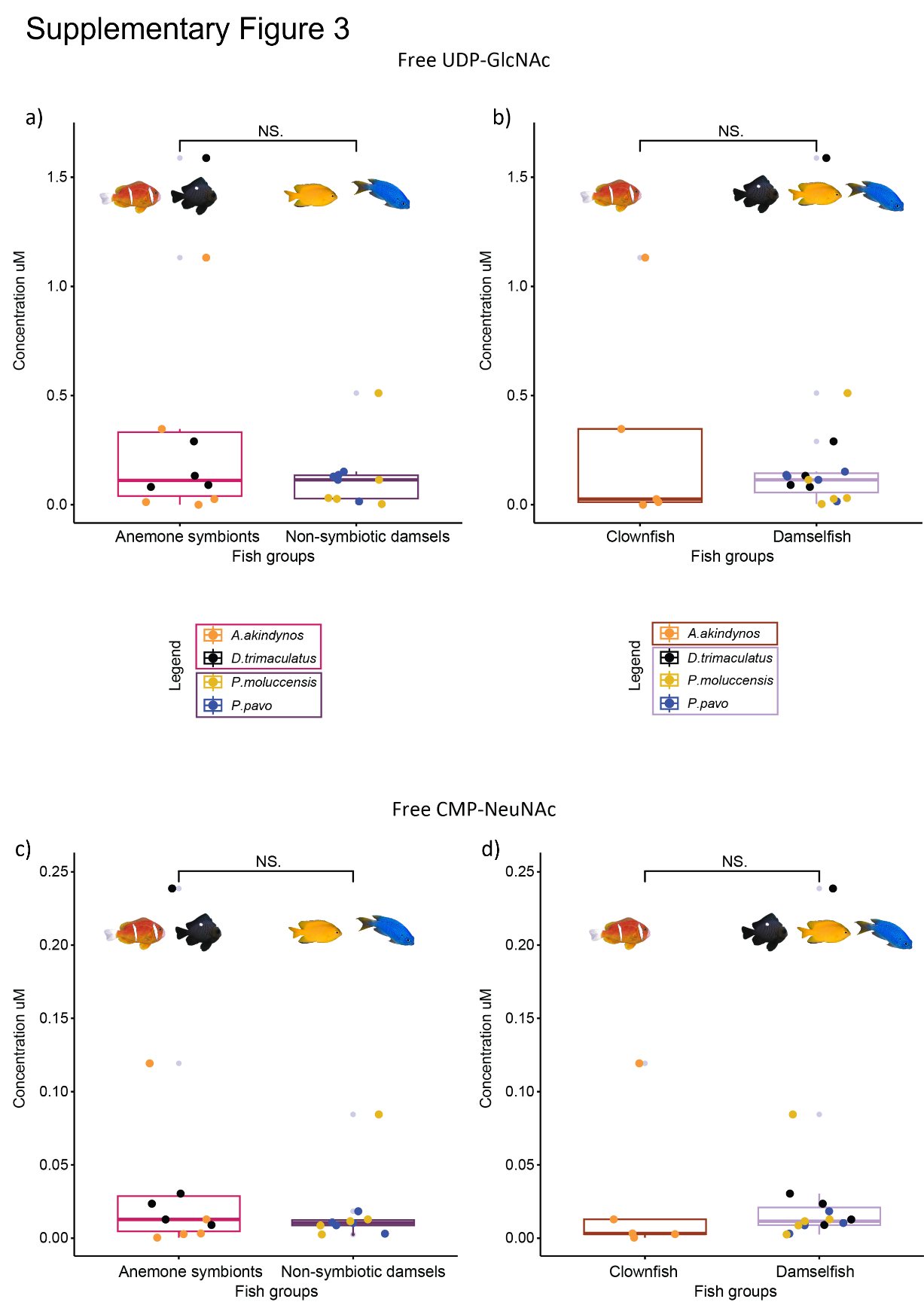

Supplementary figure 3 Boxplots of free UDP-GlcNAc (a) and (b) and free CMP-NEuNAc is represented in (c) and (d) measured in the fish species of this study (n = 5 individuals per species). The p-values above the boxplots were calculated using the anova test and contrasting anemone symbiotic fishes and non-symbiotic damselfishes, as well as the clownfish, A. akindynos, to the three other damselfish species. A. akindynos and D. trimaculatus are grouped as anemone symbionts whereas P. moluccensis and P. pavo as non-symbiotic damsels. The horizontal line in the boxplots represents the median concentration of UDP-GlcNAc and CMP-NeuNAc and the whiskers delimit the upper and lower quartiles. The grey dots neighboring a colored dot beyond the whiskers represent the outliers.
